# Emergence of clustered synapses during the development of a nervous system

**DOI:** 10.1101/2025.02.22.638574

**Authors:** Yuval Balshayi, Eduard Bokman, Alon Zaslaver

## Abstract

Synaptic organization is central for proper transmission of neural information. Studies in invertebrates and mammalian cortices show that synapses are clustered along neurite extensions, an organization that promotes key functional roles. Here we studied how these synaptic clusters emerge during the development of a nervous system. Leveraging the available *C. elegans* connectomes that span all larval developmental stages, we show that clustered synapses are formed at the early stages of the neural network development and that their occurrence further increases throughout development. These synaptic clusters significantly constitute small neural circuits that endow the network with important functional roles, such as feedback between mutually synapsing neurons and information transfer in mutually regulated neurons. Moreover, clustered synapses emerge early on during development of the head motor system, where they facilitate the crucial 3D head swings. Finally, the synaptic clusters within these key neural circuits are maintained throughout all developmental stages and are robustly found across different individuals, further accentuating their central functional roles in neural networks.

## Introduction

Neurons form polarized cell structures, known as axons and dendrites, where axonal extensions from one neuron signal dendrites of downstream neurons. In mammalian cortices, these axon-to-dendrite connections form elaborate tree-like structures in which arborized axons connect to multiple post-synaptic neurons, and each post-synaptic neuron may receive and integrate signals from multiple axons via its own tree-like dendritic structure.

It is generally accepted that axons and dendrites follow Peters’ rule, which postulates synaptic connections are randomly formed wherever axons pass in close proximity to a dendrite ^1,2^. Contemporary technologies, that extract connectome-wide structures with fine synaptic resolution, indicate that the spatial organization of synapses is not random. Instead, post-synaptic contacts exhibit a clustered organization ^3–6^. For example, in the rat barrel cortex, the vast majority of overlapping axons and dendrites are not connected, and the fraction that is connected forms synaptic clusters by connecting along the same dendritic branch ^5–7^. This clustered synaptic organization is presumably instrumental for signal integration and for accurate neural firing outputs ^8–11^. Furthermore, it is thought that such clusters play important roles in developmental and experience-dependent plasticity ^12–15^.

Interestingly, a clustered synaptic organization was also observed in the simple nervous system of *C. elegans* worms ^3^. Comprising 302 neurons, connected via ∼8000 chemical synapses and ∼1000 gap junctions, the *C. elegans* neural network is the first fully-mapped connectome for which the positions of all chemical and electrical connections are available ^16–19^. In particular, this clustered synaptic organization was demonstrated within the *C. elegans* neuropil, known as the nerve ring, where most head neurons synapse one another in a densely packed manner ^16,20,21^. In contrast to the elaborate tree-like structures of cortical neurons, the vast majority of *C. elegans* neurons are simply structured, comprising uni-or bi-polar neurites. These neurites are bundled within the nerve ring and the clustered synaptic contacts are formed along these bundled sieves.

The compact size of the *C. elegans* neural network and the simple structure of the neurons may limit the computational capacity of the network. To overcome this limited computational power, it has been suggested that the clustered synaptic organization supports local compartmentalized computations where distinct computations may be performed in parallel along a single neurite ^3^. Indeed, functional analysis of neural activity revealed neurons in which calcium dynamics were observed in local compartmentalized regions within neurites ^22–26^. For example, the RIA-type interneurons control the animals’ head bending that results from compartmentalized and reciprocal activity between the dorsal and ventral parts of its neurite ^23,24,26,27^. In the RIS neuron, local activity in one neurite branch induces locomotion stop, while co-activation together with another axonal branch promotes reversal ^22^.

Emergence of clustered synapses in a simple invertebrate nervous system as well as in mammalian brains highlights their importance to network functions. But how does the clustered organization evolve during development and maturation of the nervous system? Do such clusters form early on during development, or do they gradually emerge as new synapses are added to the network, presumably next to pre-formed synapses? Moreover, if clustered synapses endow neural circuits with crucial functions, how stereotypic is this organization when comparing isogenic individuals?

To address these fundamental questions, we leveraged the compiled *C. elegans* connectomes spanning all developmental larval stages as well as three adult-stage connectomes ^16–18^. Our analyses reveal that clustered synapses are formed early on during development and that their proportion out of the total synapses grows throughout development. They significantly constitute neural circuits that facilitate important functional roles such as feedback, integration and motor outputs. Finally, we show that synaptic clusters are preserved throughout development and across individuals, further highlighting their potential functional roles in neural networks.

## Results

### Clustered synapses are formed early on and their fraction continues to grow throughout development

To study how synaptic clusters emerge throughout the development of a nervous system, we analyzed the nine available connectomes that span the different developmental stages of *C. elegans* ^16–18^. These data comprise fully mapped connectomes of four L1-stage animals (0, 5, 8, and 16 hrs post-hatching), one each of L2-and L3-stage animals (23 and 27 hrs post-hatch, respectively), and three adult animals (50 hrs post-hatch).

We began by analyzing the simplest synaptic cluster organization in which a single pre-synaptic neuron synapses onto a post-synaptic neuron (**Figure 1A**). We defined synapses to be clustered if the distance between the actual position of the synapses along the neurites is significantly shorter than the distance of a random set of synaptic positions. Importantly, the position of the randomly placed synapses is drawn from proximal neurite regions, regions that are sufficiently close to one another and in which functional synapses could, in theory, be formed ^3^ (see Methods). For example, in the young adult animal, the DVC and AVAL neurons share a proximal stretch of neurites (**Figure 1B**). Synapses between these neurons could be randomly distributed along this proximal stretch, and yet, they are significantly clustered (p<0.001, permutation test) within a confined region.

**Figure 1.**
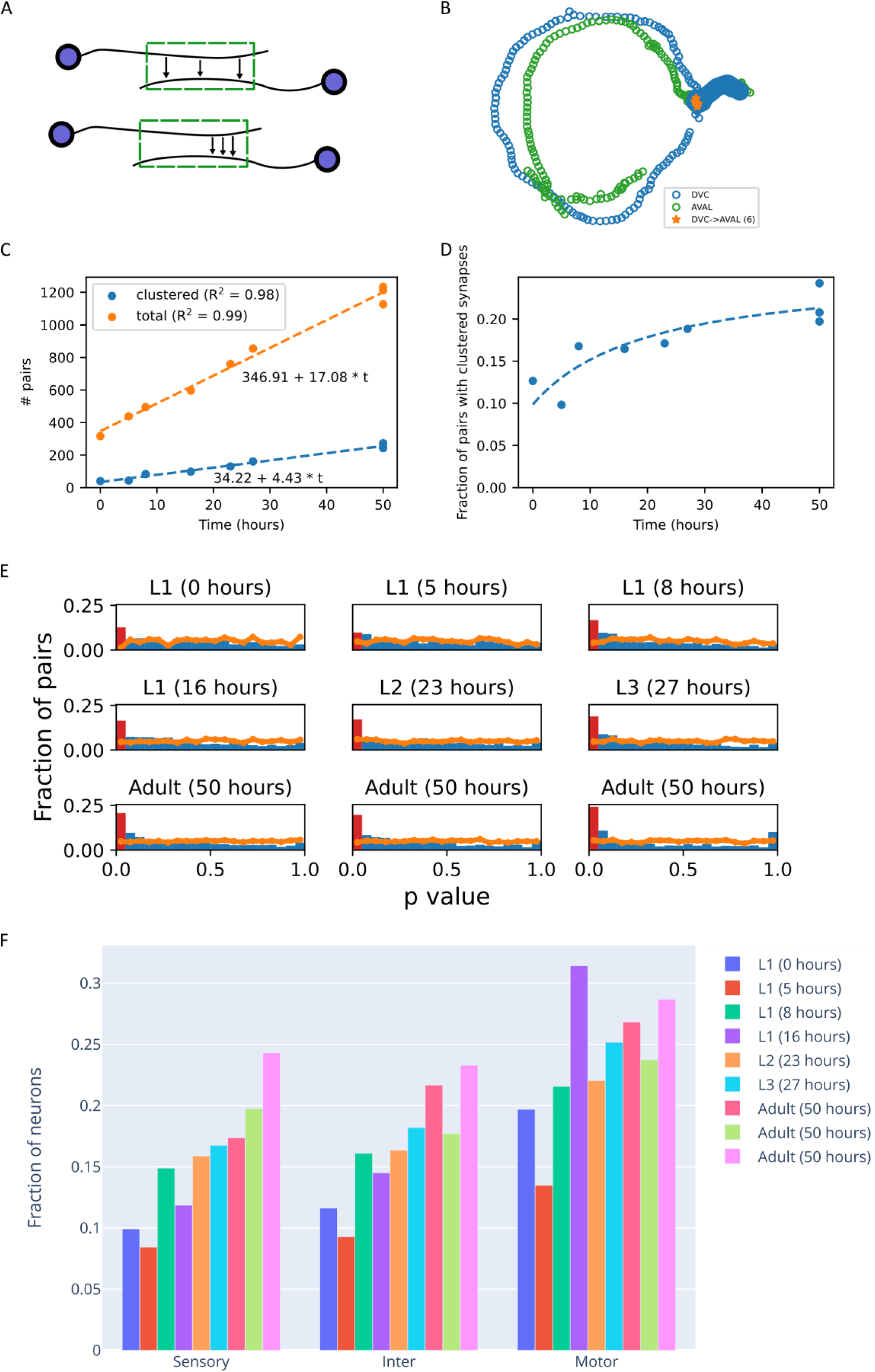
Clustered synapses are formed early during development and their fraction continues to grow throughout development. (A) Synapses may be randomly distributed (top) or clustered in a specific region (bottom) along proximal neurite regions, regions that are sufficiently close to form a synapse (green box). (B) An example of clustered synapses (orange, p<0.001) along the proximal neurite region (filled blue points) between the pre-synaptic DVC neuron and the post-synaptic AVAL neuron in a young adult animal (from dataset 8 in ^18^). (C) Number of connected pairs of neurons where a pre-synaptic neuron connects to a post-synaptic neuron. Orange: total number of connected pairs. Blue: number of connected pairs in which the synapses are clustered. (D) Fraction of connected pairs of neurons with clustered synapses out of the total number of connected pairs (trend line is the calculated from the linear fits in (C). (E) Permutation analysis shows that the fraction of connected neurons with clustered synapses is significantly enriched within the network (p<0.05), and grows throughout development. Shown are the p-value distributions when shuffling the synapses of the pre-synaptic neuron along regions of the neurite that are proximal to the post-synaptic neuron (see Methods). Orange line denotes the results of the random shuffle. Bars denote the analysis of the actual registered synapses, and a red bar shows the fraction at p<0.05. (F) Fraction of connected neural pairs with clustered synapses out of the total number of connected pairs of the same neuron type grouped according to their functional layer in the network.

During development, the total number of neural pairs connected via a synapse grows linearly, as does the number of neural pairs connected via clustered synapses (**Figure 1C**). Yet, the fraction of connected neurons sharing clustered synapses (out of the total connected pairs) increases non-linearly during development from 10% after hatch to ∼20% at the adult stage (**Figure 1D**). Notably, the fraction of pairs with clustered synapses grows during development and is significantly overrepresented throughout all developmental stages (**Figure 1E**, and see Methods for details of statistical analyses).

We next asked whether the connected pairs of neurons that form clustered synapses appear within specific layers of the network. For this, we considered the three main layers of the network, sensory, inter-, and motor neuron layers, and assigned individual neurons accordingly. The highest fraction of clustered synapses is found in the motor system neurons. This fraction is up to two-fold higher than the fraction found in neurons of the sensory and interneuron layers. Throughout development, the proportions of these clustered neurons grow similarly within each neural layer. Together, clustered synapses, abundantly found in motor neurons, are formed early on during development and their proportion among all connected neurons increases during development.

### Mutually-connected neurons form clustered synapses that appear at later developmental stages

Mutually connected neurons are small building blocks that endow neural networks with important functional roles (**Figure 2A**). These feedback circuits may serve as amplifiers or short-term memories in the case of positive feedback, and oscillators in the case of negative feedback ^28–31^. As these functional roles may be implemented by clustered synapses in a local compartmentalized manner ^3^, we next studied how such clusters emerge between mutually connected neurons throughout development (**Figure 2A-B**).

**Figure 2.**
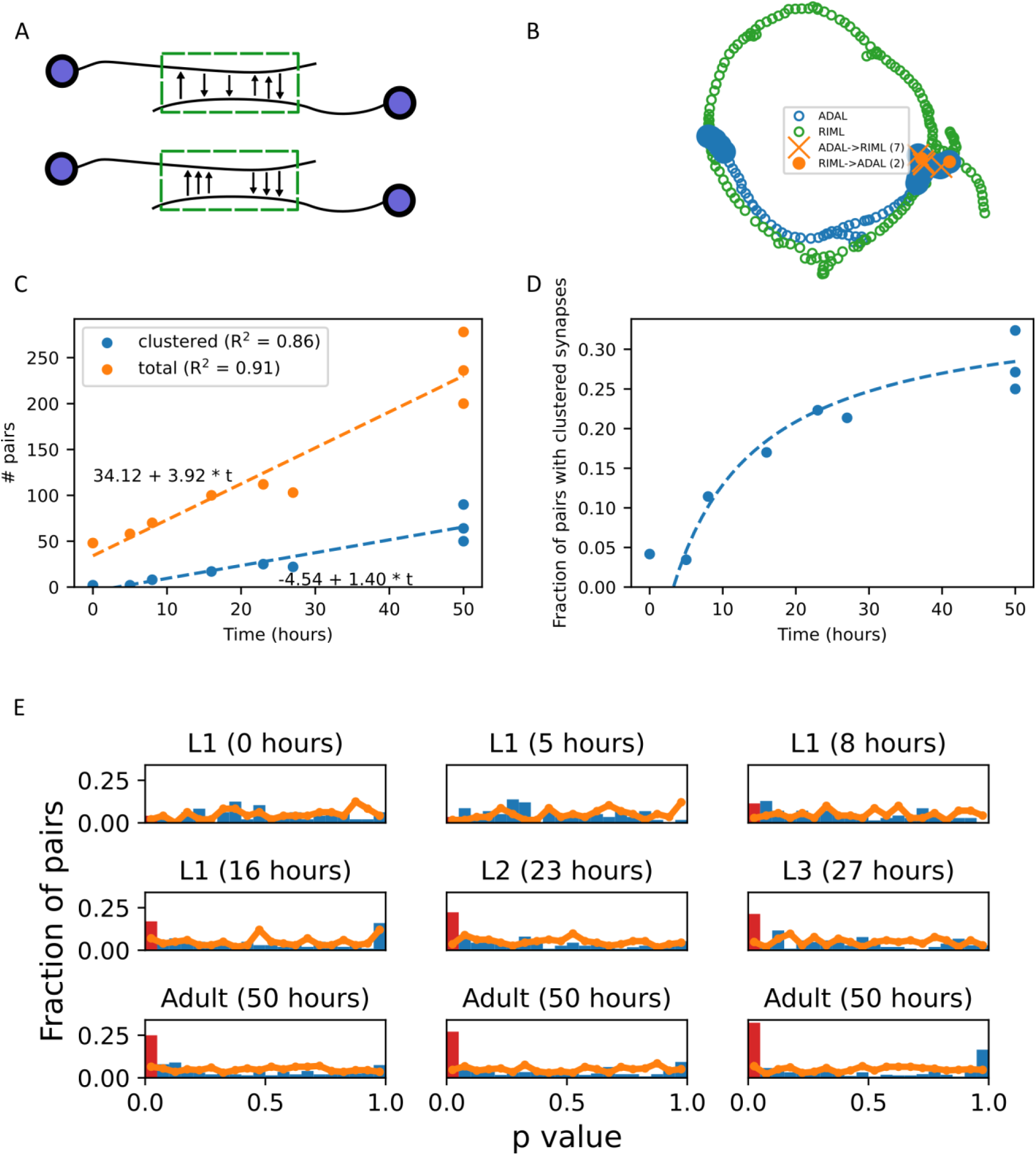
Clustered synapses between mutually-connected neurons appear at later developmental stages. (A) Illustration of possible synaptic organization between mutually synapsing neurons. Top: synapses are randomly distributed along proximal regions of the neurites. Bottom: ingoing and outgoing synapses are segregated and each forms clustered organization along proximal regions of the neurites. (B) An example of mutually synapsing neurons sharing clustered synapses. Blue and green circles denote the neurite skeletons of ADAL and RIML, respectively. Filled blue circles show their proximal neurite regions. Orange X: position of the synapses from ADAL to RIML. Orange circle: position of the synapses from RIML to ADAL. (C) Total number of mutually synapsing neurons. Blue: number of mutually synapsing neurons in which the synapses are clustered. (D) Fraction of mutually synapsing neurons showing clustered synaptic layout out of the total number of mutually synapsing neurons. (E) Bootstrap analysis showing that the fraction of connected neurons with clustered synapses grows throughout development. Shown are the p-value distributions when shuffling the positions of the synapses between the mutually synapsing neurons along the proximal regions.

In general, both the total number of mutually connected neurons as well as the number of mutually connected neurons that share clustered synapses grow linearly during development (**Figure 2C**). However, the fraction of connected pairs with clustered synapses increases significantly more from ∼5% at the early L1 stage to ∼25% at the adult stage (**Figure 2D**). Moreover, while the fraction of mutually connected neurons with clustered synapses is not significantly enriched in the connectome at the L1 stage, this fraction grows quickly to become significantly enriched from the L2 stage and onward (**Figure 2E**). These findings suggest that clustered synapses between mutually-synapsing neurons may endow the network with pivotal functional roles, yet these functions become particularly important in the later stages of the developing nervous system.

### Tri-neuron circuits with clustered synapses form early during development to control head movement via the RIA interneurons

We next studied how synaptic clusters develop within neural circuits made of three neurons. We focused on simple circuits that lack feedback, in which two neurons connect with a third neuron as either pre-or post-synaptic neurons and regardless of other interconnections (**Figure 3A**). These circuits can be classified as: **(I)** two neurons that synapse onto a third shared post-synaptic neuron, forming an input function known as mutually regulating neurons; **(II)** a single neuron that regulates two downstream post-synaptic neurons, a layout known as mutually regulated neurons; **(III)** a linear path of information flow in which one neuron synapses a second neuron, which in turn synapses a third neuron. In all these layouts, clustered synapses may be formed on the neurite of the shared neuron, and by this, support local compartmentalized activity (**Figure 3B**).

**Figure 3.**
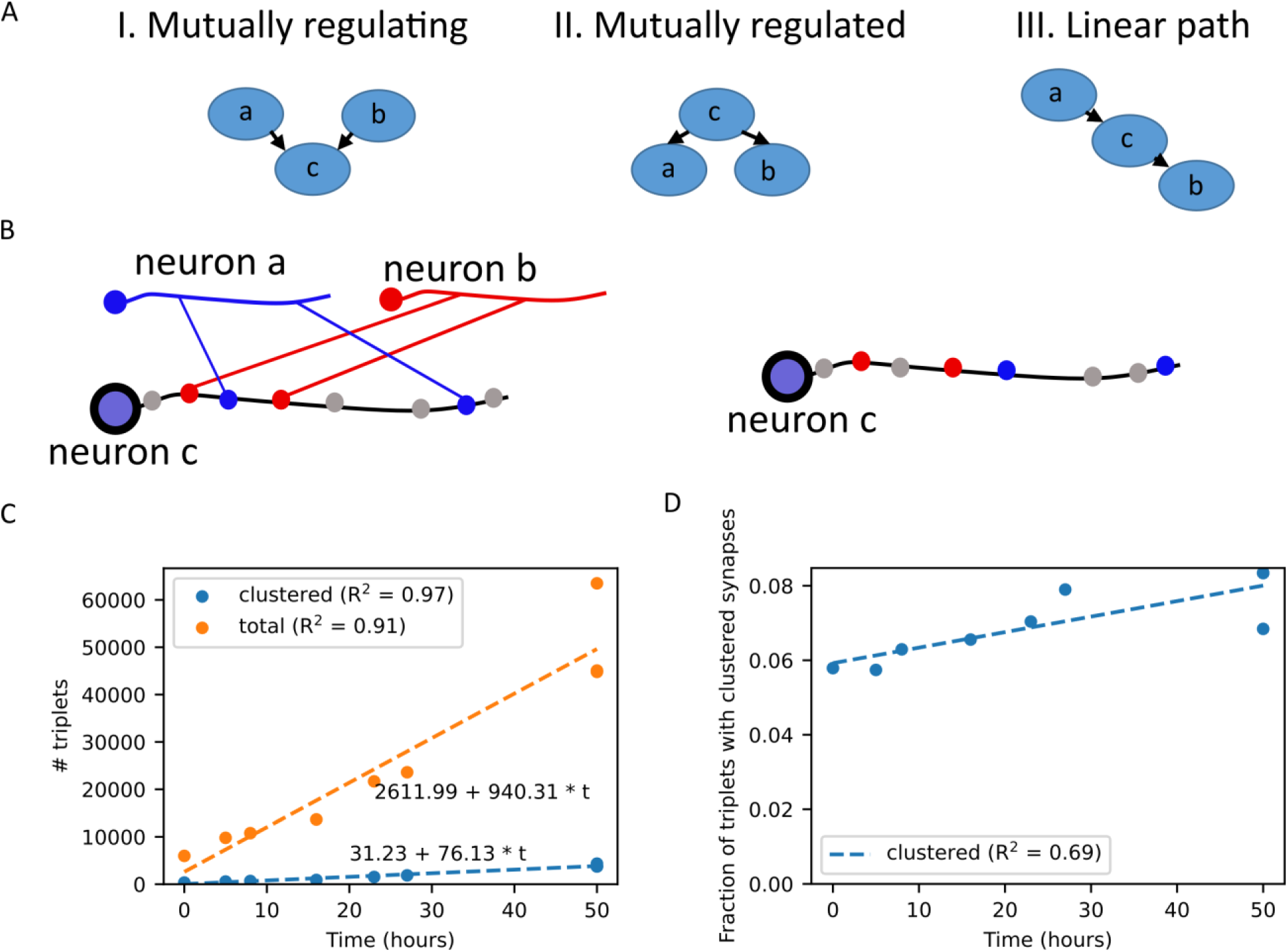
Emergence of clustered synapses in circuits consisting of three neurons. (A) Layouts of simple three-neuron circuits in which local compartmentalized activity may be performed on the neurite of the shared neuron: (I) Mutually regulating circuit: a shared neuron {c} is post-synaptic to the two pre-synaptic neurons {a} and {b}; (II) Mutually regulated circuit: a shared pre-synaptic neuron {c} synapses onto two post-synaptic neurons {a} and {b}; (III) A linear circuit: the shared neuron {c} is post-synaptic to neuron {a} and pre-synaptic to neuron {b}. (B) Illustration of a mutually regulating circuit (layout (I) of panel A) and the analysis of the clustered synaptic organization on the mutual neuron {c}. We compared the mean of the distances between the actual registered synapses of neurons {a} and {b} along the neurite of the common neuron {c} (left panel) to the distances between the synaptic positions of one neuron (neuron {b} marked with red synapses, in this case) and randomly selected synaptic positions along the proximal regions on neurite {c} (right panel). (C) Number of tri-neuron circuits. Orange: total number of circuits. Blue: number of circuits in which the synapses are clustered. (D) Fraction of tri-neuron circuits showing clustered synaptic layout out of the total number of tri-neuron circuits.

The number of such tri-neuron circuits that share clustered synapses mildly grows when compared to the total number of such tri-neuron circuits (**Figure 3C**). In fact, the fraction of these tri-neuron circuits with clustered synapses grows from just 6% at the L1 stage to 8% at the adult stage (**Figure 3D**). Thus, the vast majority of the clustered tri-neuron circuits are formed early on during development. As such, we hypothesized that these circuits fulfill critical functional roles throughout all life stages.

We therefore analyzed whether these tri-neuron circuits are associated with a specific type of the tri-neuron circuits while additionally considering the neuron layers comprising the circuit, a feature that may indicate their potential functional roles. For this, we assigned neurons to one of the network layers: sensory, inter-, and motor neuron layers. Overall, there are 63 different configurations to generate the three circuit layouts while considering the three neuron types. Notably, of the 63 possible configurations, only four circuits were overrepresented within the network throughout most developmental stages (**Figure 4A-B, Supplementary figure 1A**).

**Figure 4.**
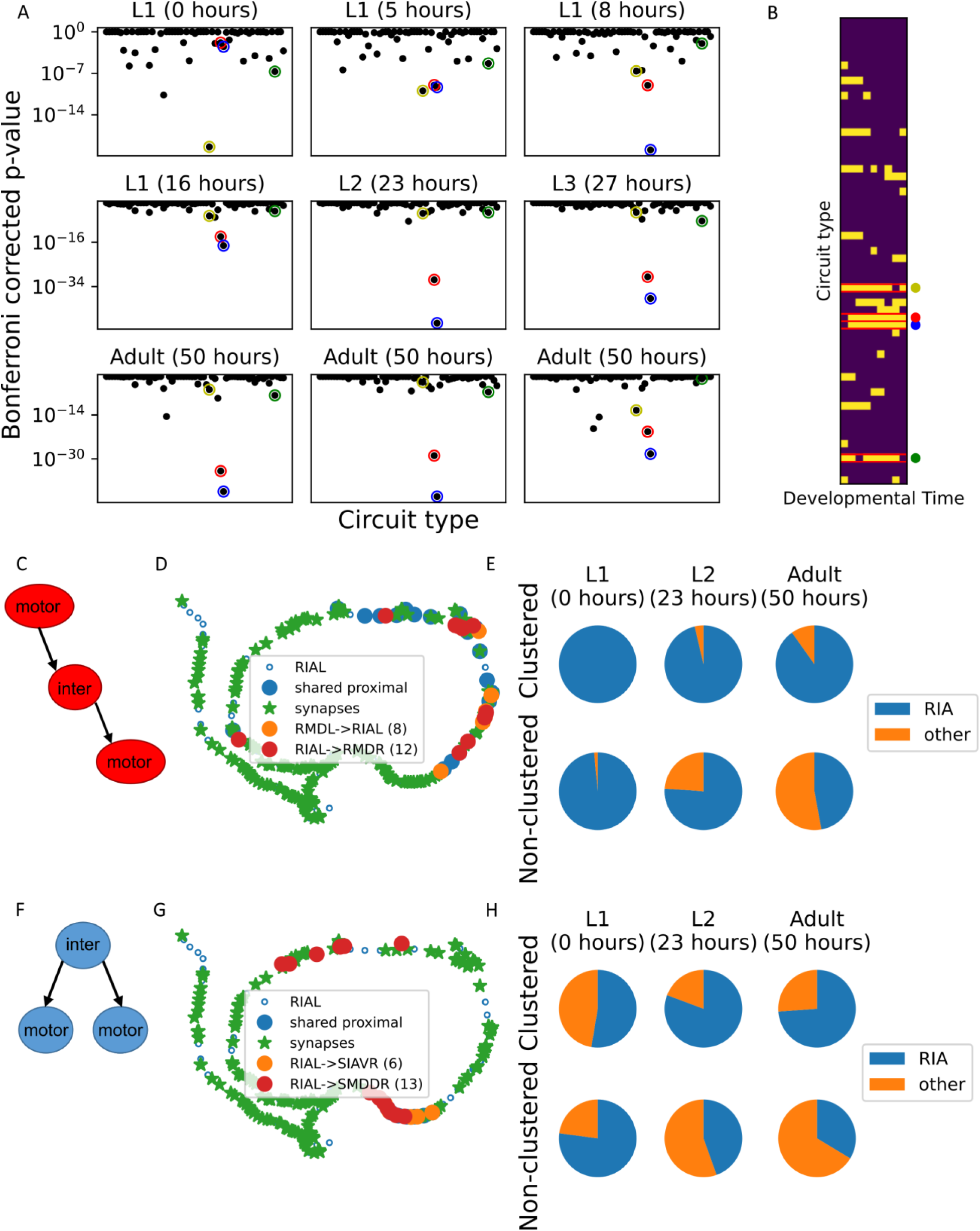
Overrepresented tri-neuron circuits are enriched with the RIA interneurons sharing clustered synapses with motor neurons. (A) Significance of the occurrence of the 63 different tri-neuron circuits with clustered synapses comprising sensory, inter-, and motor neurons (see also methods). (B) Heat map showing which circuit types are significantly clustered (yellow) in each dataset (hypergeometric test). Four circuit types that stood out are marked by colored circles: (Yellow) a presynaptic interneuron which connects to two postsynaptic interneurons; (Red) an interneuron which is both pre-and postsynaptic to motor neurons; (Blue) a presynaptic interneuron which connects to two postsynaptic motor neurons; (Green) a motor neuron which is presynaptic to both an interneuron and a motor neuron. (C) A schematic view of a linear transmission circuit in which an interneuron is both pre-and post-synaptic to two motor neurons. (D) An example of a linear transmission circuit with clustered synapses. Synapses from RMDL and to RMDR motor neurons are clustered (p<0.001) along the neurite of the RIAL interneuron (taken from the L2 developmental stage). (E) Fractions of the overrepresented linear circuits with clustered (top) and non-clustered (bottom) synapses with RIA as the connecting interneuron. Shown are three representative time points along the developmental axis (see **Supplementary figure 2A** for analysis of all developmental stages). (F) A schematic view of a mutually regulating circuit in which an interneuron synapses onto two postsynaptic motor neurons. (G) An example of a mutually regulating circuit with clustered synapses. Synapses to the SIAVR and SMDDR motor neurons are clustered (p=0.001) along the neurite of the presynaptic interneuron RIAL (taken from the L2 developmental stage). (H) Fractions of the overrepresented mutually regulating circuits with clustered (top) and non-clustered (bottom) synapses with RIA as the shared pre-synaptic interneuron. Shown are three representative time points along the developmental axis (see **Supplementary figure 2B** for analysis of all developmental stages).

Interestingly, a detailed analysis revealed that the RIA interneuron type is significantly overrepresented in two of these circuits throughout all developmental stages (except for the very early L1 stage, **Supplementary figure 1B**). The first circuit depicts a linear layout in which the interneuron is pre-and postsynaptic to motor neurons (**Figure 4C-D**). The vast majority of these circuits, which also form clustered synapses, consist of RIA as the interneuron throughout all developmental stages (**Figure 4E**). In contrast, in non–clustered synapses, the fraction of RIA drops during development to be half of all interneurons at the adult stage (**Figure 4E and Supplementary figure 2A**). In the second circuit, the RIA interneuron is presynaptic to two motor neurons (**Figure 4F-G**). Of the circuits with clustered synapses, the fraction of RIA is ∼half at the L1 stage and this fraction grows to ∼75% at the adult stage (**Figure 4H and Supplementary figure 2B**). In contrast, the fraction of RIA in circuits with non-clustered synapses ∼75% at the early L1 stage and this fraction drops to ∼30% at the adult stage.

These findings indicate that only a few tri-neuron circuits are overrepresented having clustered synapses throughout most developmental stages. Moreover, the pair RIA interneurons are particularly enriched within two of the circuits where they either receive or transmit information to motor neurons. Indeed, the RIA neurons are central in regulating head movement via compartmentalized activity ^23,24,27^, and their significant overrepresentation in forming clustered synapses with head motor neurons throughout all developmental stages suggests a central role for these clustered synapses in mediating the finely-controlled head movements.

### Clustered synapses are conserved in the developing nervous system

The above analyses showed that clustered synapses are formed early on during development and that they appear preferentially in specific circuits. But how variable are the identities of the clustered synapses between individuals and how well preserved are they across the different developmental stages?

To address these questions, we performed a pairwise analysis across all developmental stages and calculated the % of shared neural pairs with clustered synapses (out of the total pairs with clustered synapses between each two developmental stages (**Figure 5A top-left**, and see Methods for details). The mean percent of shared clustered synapses is 36%±10.5 when comparing across all developmental stages, and this mean rises to 41%±6.8 when comparing between sequential developmental stages. Furthermore, comparing connectomes of adults only reveals a similar ∼40% preservation degree of clustered synapses. When repeating this analysis for neural pairs connected via non-clustered synapses, a higher fraction (84.6%±5.2) of shared neural pairs is obtained (**Figure 5 top-right**). However, neural pairs connected via non-clustered synapses constitute the majority of connected neural pairs: In fact, non-clustered synapses make 80%-90% of all neural connections across the different developmental stages, while clustered synapses constitute only 10%-20% of the total neural connections (**Figure 1D**). It is therefore expected that non-clustered neural pairs will show a higher preservation degree (see Methods for extended explanation). Surprisingly however, when controlling for the number of neural pairs in each group, we find that the expected % of shared clustered synapses is significantly lower (17.1%±4.2, **Figure 5A middle panels**). In contrast, the mean of the expected fractions of non-clustered synapses remains similar to the non-randomized pairs (∼82.0%±4.8). Thus, the fraction of shared clustered synapses is significantly higher than randomly expected, as also demonstrated by the z-score analysis, where the mean Z-score values for clustered and the non-clustered synapses is 5.0±2.7 and 1.8±1.7, respectively (**Figure 5A bottom panels**). Particular high preservation degree of clustered synapses was observed from the L2 stage and on, indicating that clustered connections formed at the L2 stage are more prone to remain through subsequent developmental stages.

**Figure 5.**
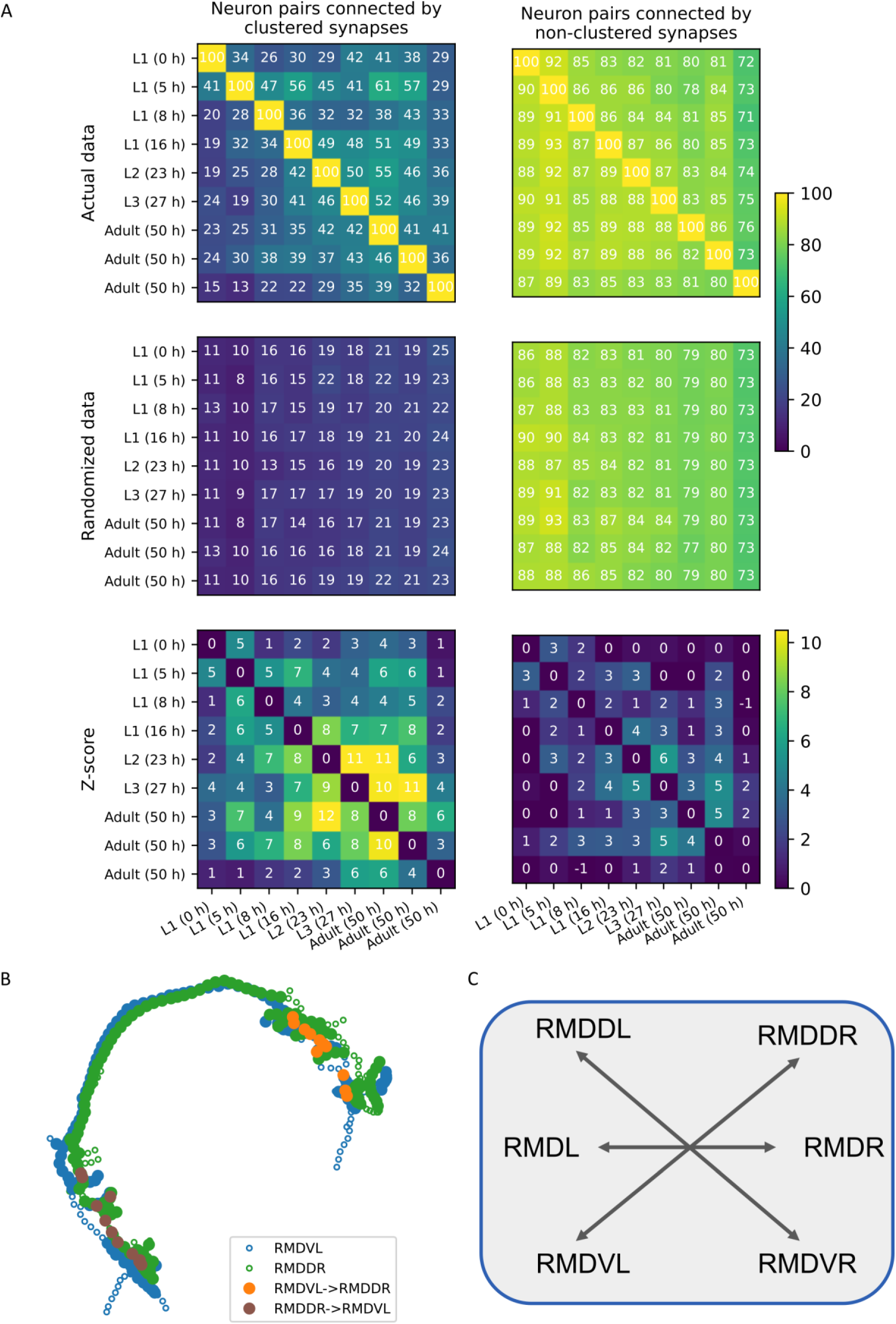
Clustered synapses are significantly preserved throughout development and across individuals. (A) Top: A pairwise analysis indicating the % of neural pairs with clustered synapses in one developmental stage (denoted by the row) that also have clustered synapses in the other developmental stage (denoted by the column). Middle: Repeating the analysis using bootstrap to correct for the number of neural pairs with clustered and non-clustered synapses (see Methods). N=50 iterations. Bottom: Z-scores calculated based on the top and the middle matrices. Left and right panels denote the neural pairs connected by clustered and non-clustered synapses, respectively. The values of the right most and the bottom are for the adult animal connectome compiled by ^17^. All developmental stages as well as the two other adult connectomes are from ^18^. Colorbars indicate percent values for the top and the middle matrices, and Z scores for the bottom matrices. (B) The neurites between the RMD neurons form clustered synapses. Shown as example the synapses along the neurites of the RMDVL (blue) and the RMDDR (green) neurons. These clustered synapses are preserved throughout all developmental stages. Filled and open circles indicate proximal and non-proximal regions of the neurites, respectively. Orange: positions of synapses from the RMDVL to the RMDDR neurons. Brown: synapses in the opposite direction. (C) A schematic layout of the contralateral connectivity of the RMD neurons. These contralateral neurons share clustered synapses throughout all developmental stages.

The compiled connectomes show considerable inherent variability with regard to the number of synapses registered between pairs of connected neurons ^18^. Consequently, neural pairs sharing one or two synapses in one connectome may lack these synapses in another connectome, and hence, not be categorized as connected. This variability, stemming from the ‘low numbers’ principle, may therefore bias our analysis. To control for this inherent variability, we repeated the analysis, this time considering sets of connected pairs that share at least three or four synapses. Similar higher values of shared clustered synapses were obserbed in these more stringent analyses, further underscoring the notion that clustered synapses are significantly more conserved than non-clustered synapses (**Supplementary figures 4-5**).

These results indicate that while the overall preservation of clustered synapses is merely ∼40%, this preserved fraction is significantly higher than the preservation obtained for non–clustered synapses. Furthermore, when comparing the preservation degree of clustered synapses of the three adult-stage connectomes only, a similar % preservation is observed, suggesting that this may be the upper limit for preservation detection within a population of isogenic animals.

Interestingly, a single type of neuron, RMD, is consistently found to be connected via clustered synapses throughout all developmental stages (**Figure 5B** and **Table 1**). The RMD type includes six motor neurons with left/right and dorsal/ventral symmetry. These neurons form clustered synaptic connections contra laterally (for example, the dorsal-right RMDDR neuron and the ventral-left RMDVL neuron, **Figure 5C**), and each neuron also connects to head muscles via neuromuscular junctions. This way, the RMD neurons control head movement in all directions and the clustered contralateral synapses presumably allow efficient synchronization when controlling opposite side muscles. Furthermore, the RIA type neurons, which are the main neurons to participate in clustered synaptic contacts (**Figure 4**), also form bi-directional clustered synaptic contacts with the RMD neurons to control head movement. These bidirectional clustered synapses are also preserved throughout development from the middle L1 stage and on (**Table 1**). Together, these findings demonstrate how crucial functional roles are mediated via clustered synaptic structures which evolve in the network early on during development and are preserved to the adult stage. Moreover, they are non-variably appear in all three adult animals, further underscoring their importance in controlling central locomotion functions.

**Table 1.** Synaptic clusters preserved throughout development. The number of synapses between pairs of neurons that share clustered synapses throughout the different developmental stages. The connections between all pairs of neurons have significant clustered synapses, except for those indicated as not significant (ns). Note the top rows consisting of the RMD neurons which are interconnected via clustered synapses throughout all developmental stages.

| Pre | Post | L1<br>(0 h) | L1<br>(5 h) | L1<br>(8 h) | L1<br>(16 h) | L2<br>(23 h) | L3<br>(27 h) | YA<br>(50 h) | YA<br>(50 h) |
| --- | --- | --- | --- | --- | --- | --- | --- | --- | --- |
| RMDDL | RMDVR | 4 | 6 | 6 | 3 | 9 | 6 | 8 | 11 |
| RMDDR | RMDVL | 4 | 5 | 6 | 5 | 9 | 10 | 14 | 12 |
| RMDL | RMDR | 2 | 4 | 5 | 8 | 7 | 7 | 7 | 12 |
| RMDVL | RMDDR | 4 | 5 | 6 | 5 | 10 | 9 | 16 | 11 |
| RMDVR | RMDDL | 3 (ns) | 6 | 5 | 6 | 10 | 11 | 14 | 10 |
| AIBL | SMDDR | 2 (ns) | 2 (ns) | 3 | 3 | 3 | 5 | 4 | 5 |
| AIBR | SMDDL | 2 (ns) | 4 | 3 | 2 | 5 | 4 | 2 (ns) | 6 |
| RMDL | RIAR | 2 (ns) | 2 (ns) | 4 | 3 | 5 | 4 | 8 | 8 |
| RMDR | RIAL | 2 (ns) | 3 (ns) | 3 | 3 | 5 | 6 | 5 | 7 |
| RMDR | RMDL | 2 (ns) | 3 | 7 | 6 | 9 | 8 | 5 | 7 (ns) |
| SAADL | RIML | 2 | 2 | 3 (ns) | 7 | 3 | 5 (ns) | 8 | 5 |
| SAADL | RIMR | 2 | 2 (ns) | 2 (ns) | 4 | 6 | 3 | 9 | 10 |
| SAAVL | AVAL | 0 (ns) | 4 | 3 (ns) | 5 | 11 | 10 | 19 | 19 |
| SAAVR | AVAR | 3 (ns) | 5 | 3 (ns) | 4 | 13 | 10 | 26 | 21 |

## Discussion

Rapidly advancing technologies now make it possible to document individual synapses within intact brain regions and to compile detailed synapse-based connectomes, revealing that synapses are organized in clusters ^4–6,17–19^. Synaptic clustering is presumably shared across the animal kingdom as clustering has been observed in mice cortices as well as in the invertebrate *C. elegans* worm. This converging structural feature strongly suggests that synaptic clusters have key functional roles in neural circuits that allow signals to propagate in an accurate and timely manner ^10^. Herein, leveraging the available time-series connectomes of *C. elegans* ^16–18^, we studied how these clustered synaptic structures emerge during the development of a neural network.

Naively, there are two possible scenarios for the emergence of clustered synaptic structures: These structures could be formed early on during development and maintained throughout development. Alternatively, these structures could emerge during development where initially synapses are sparsely dispersed between connecting neurons and during development new synapses are preferentially added next to pre-existing synapses to form clustered structures. Our analyses show that both processes take place in the *C. elegans* connectome. We find that key clustered synapses emerge early on during development of the nervous system and that the fraction of clustered synapses out of the total new synaptic connections in the network also grows till the adult stage (**Figure 1**).

Analysis of the *C. elegans* connectome suggests that it follows the Peter’s rule, where synapses form merely because two neurites are sufficiently close to one another ^32,33^. Our study shows that not only do such synapses form between adjacent neurites, but that a significant fraction of these synapses emerges in clusters early on during development and within small circuits that endow the network with key functions. For example, a significant fraction of the mutually synapsing neurons forms clustered synapses (**Figure 2**). These bidirectional connections that form feedback loops are thought to provide the network with key functions such as oscillations and amplifiers in case of negative and positive feedback loops, respectively ^28,30,31^.

Analysis of small circuits composed of three neurons revealed a few layouts that stood out as significantly overrepresented throughout development to consist clustered synapses. Strikingly, the bilateral RIA interneurons appear in the vast majority of these circuits (**Figure 4**). The RIA interneurons output primarily onto motor neurons, including the RMD motor neurons that control head movement ^23,24,27^. Interestingly, the RMD neurons themselves connect to their contralateral partners via clustered synapses as well as to the RIA neurons. Moreover, all these clustered synapses are formed early on during development and are preserved throughout all developmental stages (**Figure 5 and Table 1**). Thus, emergence of clustered synapses early on during development may support key motor functions that are crucial at the very first hours of the developing animal and throughout its entire life.

The mutually connected neuron pairs and the tri-neuron circuits are embedded within a dense neural net in which each neuron is part of several different circuits. Indeed, it has been shown that mutually synapsing as well other tri-neuron circuits make small building blocks within larger circuits ^30^. These larger circuits consist of interneurons that integrate sensory information and motor neurons that receive multiple inputs from interneurons to promote a synchronized coordinated movement. Clustered synaptic structures along the neurites of these inter and motor neurons may be crucial for such fine locomotion patterns.

Notably, functional studies of selected interneurons revealed calcium transients within compartmentalized regions of the neurites ^22–26,34,35^. For example, the RIA interneurons control the animals’ head movement via compartmentalized and reciprocal activity between the dorsal and ventral parts of its neurite. Compartmentalized activity was also observed in the nerve ring segment and the axonal branch of the RIS neuron: activity in the nerve ring process alone correlates with locomotion stop and coactivity with the axonal branch promotes reversal events. These studies further highlight the functional importance of synaptic clusters, demonstrating how neural information can be transferred in a neurite-local manner without involving the entire neuron. Furthermore, as several compartments may function in parallel along a single neurite, the emergence of synaptic clusters can greatly enhance the computational capacity of the entire neural network. This feature may be particularly relevant in the early stages of a developing neural network as clustered synapses in key circuits may enhance functional repertoires despite the fact that the network is not yet fully wired and many of the neural connections are yet to be established.

To this end, the *C. elegans* neuropil (the nerve ring) is organized into four layered regions (strata) that segregate early on during embryogenesis ^21^. Interestingly, a group of ∼30 ‘rich club’ neurons ^36,37^ that share multiple synaptic partners bridge across these strata (including RIA and RIS mentioned above). Synaptic clusters along the neurites of these highly-connected neurons are partitioned across different strata, presumably to transduce information across these segregated layers.

Our analyses indicate that synaptic structures are found within key neural circuits, and that they are significantly preserved throughout development and across adult individuals. However, it is somewhat surprising that the preservation degree of these clustered structures across isogenic adult animals is only ∼40% (**Figure 5**). Even when exclusively considering synaptic partners that share more than a few synapses, thus overcoming the inherent variability associated with small number of synapses, the preserved degree between adult animals reaches to ∼50% only (**Supplementary Figure 5)**. Indeed, network analyses throughout all developmental stages, including the three adult-stage connectomes, demonstrated considerable differences in both the number of synapses and in the adjacency of the neurites ^18,33^. Thus, the low preservation degree in the identity of the clustered synapses may be a direct result of this high inter-individual variability. Nevertheless, our analyses indicate that in crucial circuits, the clustered structures are preserved during development and across individuals.

Taken together, this study demonstrates that key functions are mediated via clustered synaptic structures that emerge early on during the development of a neural network. This may explain why these clustered structures are ubiquitously found in diverse animal species.

## Methods

### Datasets

Datasets used in this study were retrieved and compiled from ^3,18,19^. Each dataset was compiled from a series of electron micrographs and provides coordinates of points along the skeleton of the neurons, as well as the spatial positions of the synapses (by associating the pre-and post-synaptic positions with the skeleton points). In regions in which there were large gaps between skeleton points (defined as larger than the distance between three consecutive electron micrographs), additional points were added by interpolation so that the gaps would be approximately the slice distance.

### Testing significance of clustering of synapses between a pair of neurons

We used a permutation test (**Figures 1E and 3E**) to test whether synapses are clustered between a pair of pre-and post-synaptic neurons: the mean distance between the real synaptic positions along the pre-synaptic neuron was compared to the mean distance between randomly selected positions along the region of the neurite that is sufficiently close (proximal) to form a synapse with the post-synaptic neuron.

To determine the axonal proximal regions (the regions that could potentially form a synapse), we first calculated the median distance between all pre-and post-synaptic positions in the dataset. We then defined proximal regions as points along the presynaptic neuron that are closer to a post-synaptic neuron point than the calculated median distance (as also defined in ^3^). The permutation test was run 1000 times or for all possible permutations if there were fewer possible permutations.

The synapse clustering p-value was defined as the fraction of tests in which the mean distance between the randomly positioned synapses was less than the mean distance between the actual synapses. This test was performed for every pair of neurons connected by at least two synapses, as well as for mutually connected neurons.

### Testing clustering significance of synapses of two neurons along the neurite of a third common neuron

To determine whether the synapses of two neurons (A and B) are clustered along the neurite of a third common neuron (C), we applied the following permutation test: in each permutation, the mean distance between every pair of A and B synapses along the neurite of neuron C was compared to the mean distance between the positions of the real synapses of one of the neurons and randomly selected synapses from all synapse positions along neurite C. This test was run 1000 times while shuffling the synapses of A, and 1000 times shuffling the synapses of B. When the number of possible permutations was less than 1000, all possible permutations were considered. From each set of tests, a p-value was calculated as the fraction of tests in which the mean distance between the randomly positioned synapses was less than the mean distance between the real synapses. Finally, the p-value for clustering in the triplet was defined as the maximum between p-values when the synapses of A were shuffled and when the synapses of B were shuffled.

### Tightly-clustered tri-neuron circuits

We performed both permutation tests and hypergeometric tests to study which specific three-neuron circuits have a higher probability of forming clustered synapses along the neurite of the common neuron. For the permutation tests, we shuffled which of the tri-neuron circuits are clustered 3000 times, and compared the number of tri-neuron circuits which were clustered after shuffling with the real number of circuits that have clustered synapses. The hypergeometric tests were performed using the total number of circuits, the total number of circuits with clustered synapses, the total number of circuits of each type and the number of circuits of the specific type that have clustered synapses.

### Analysis of synaptic clusters preservation

We performed a pairwise comparison across all datasets. First, we compiled a list of all connected pairs of neurons that share at least two synapses (clustered and non-clustered) as well as a list of only the pairs of neurons that share clustered synapses (sharing at least two synapses) in each dataset. To calculate the fraction of neural pairs with clustered synapses in one dataset that are also clustered in another dataset, we counted the number of pairs with clustered synapses that appear in both lists of pairs with clustered synapses (one for each dataset) and divided this number by the number of neural pairs that appear both in the list of pairs with clustered synapses for one of the datasets and in the list of connected pairs for the other dataset, thus controlling for pairs that could not be analyzed in the second dataset, since they were not connected (**Supplementary figure 3**). For the comparison of pairs with non-clustered synapses, we repeated this analysis using the complement of the list of pairs with clustered synapses in each dataset.

To calculate the expected percentages of shared pairs between datasets, we ran the above calculation 50 times, each time shuffling the identities of the clustered pairs in one of the datasets. We then calculated the mean and standard deviation for each pair of datasets and used these values to calculate the z-score.

## Acknowledgements

We thank Cheryl Berkowitz for critical reading and valuable feedback on the manuscript draft. We also thank Rotem Ruach for the insightful advices.

## Availability of Data and Materials

All the data and the scripts that analyzed the data to produce the results and the figures herein are available through Github: https://github.com/zaslab/ce-synaptic-clustering.

## Author Contributions

AZ and YB conceived the project. YB developed the analytic tools and performed the data analyses. EB contributed statistical tools; AZ and YB wrote the manuscript.

## Supplementary figures

**Supplementary Figure 1.**
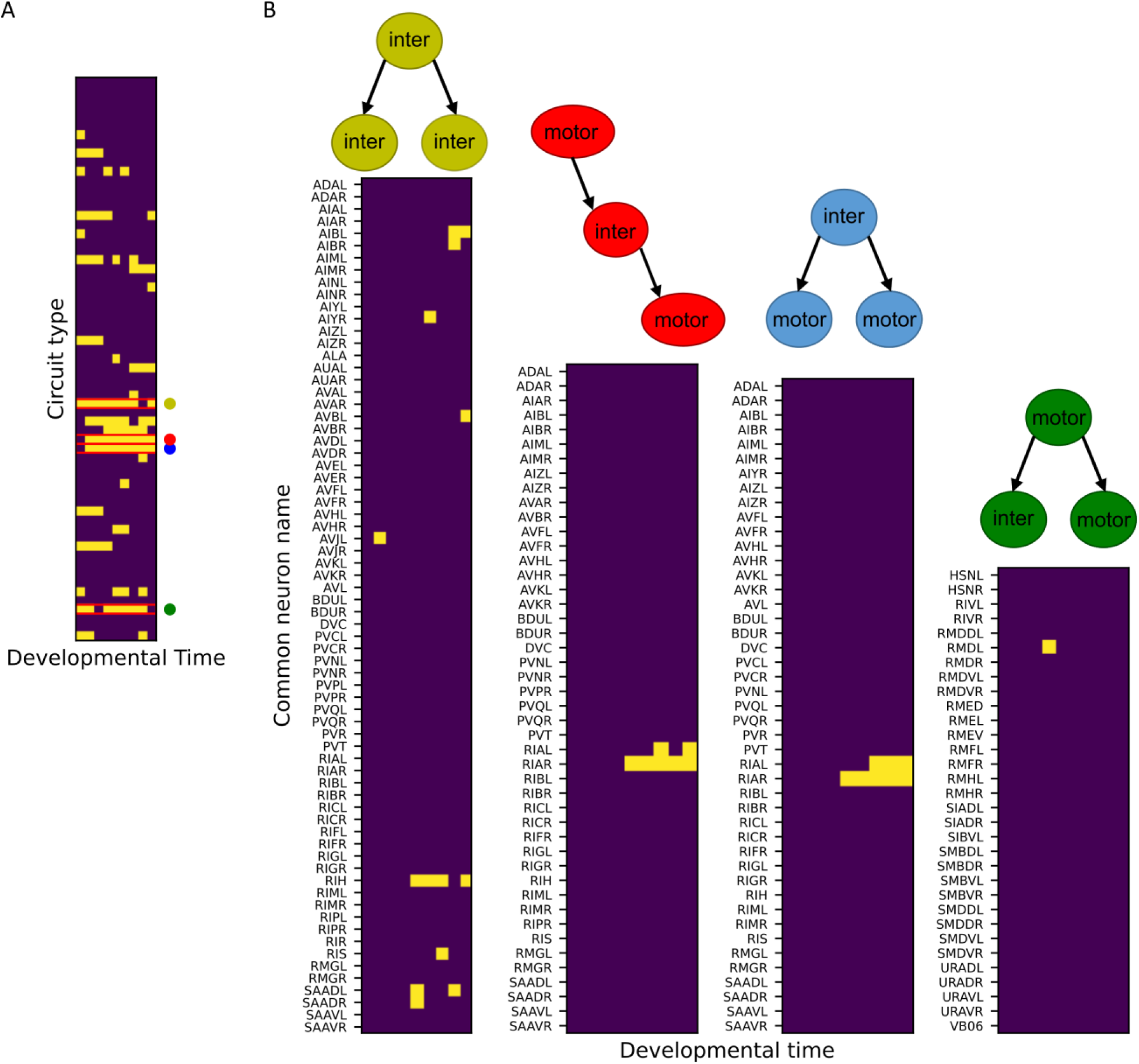
Emergence of tri-neuron circuits during development. (A) Heat map showing which circuit types are significantly clustered (yellow) in each dataset (permutation test). Four circuit types stood out, from top to bottom: a presynaptic interneuron which connects to two postsynaptic interneurons (yellow dot), an interneuron which is both pre-and postsynaptic to motor neurons (red dot), a presynaptic interneuron which connects to two postsynaptic motor neurons (blue dot), and a motor neuron which is presynaptic to both an interneuron and a motor neuron (green dot). (B) Heat map showing which neurons are over represented as having clustered synapses in each of the overrepresented clustered circuit types (left to right: a presynaptic interneuron which connects to two postsynaptic interneurons (yellow dot), an interneuron which is both pre-and postsynaptic to motor neurons (red dot), a presynaptic interneuron which connects to two postsynaptic motor neurons (blue dot), and a motor neuron which is presynaptic to both an interneuron and a motor neuron (green dot)).

**Supplementary Figure 2.**
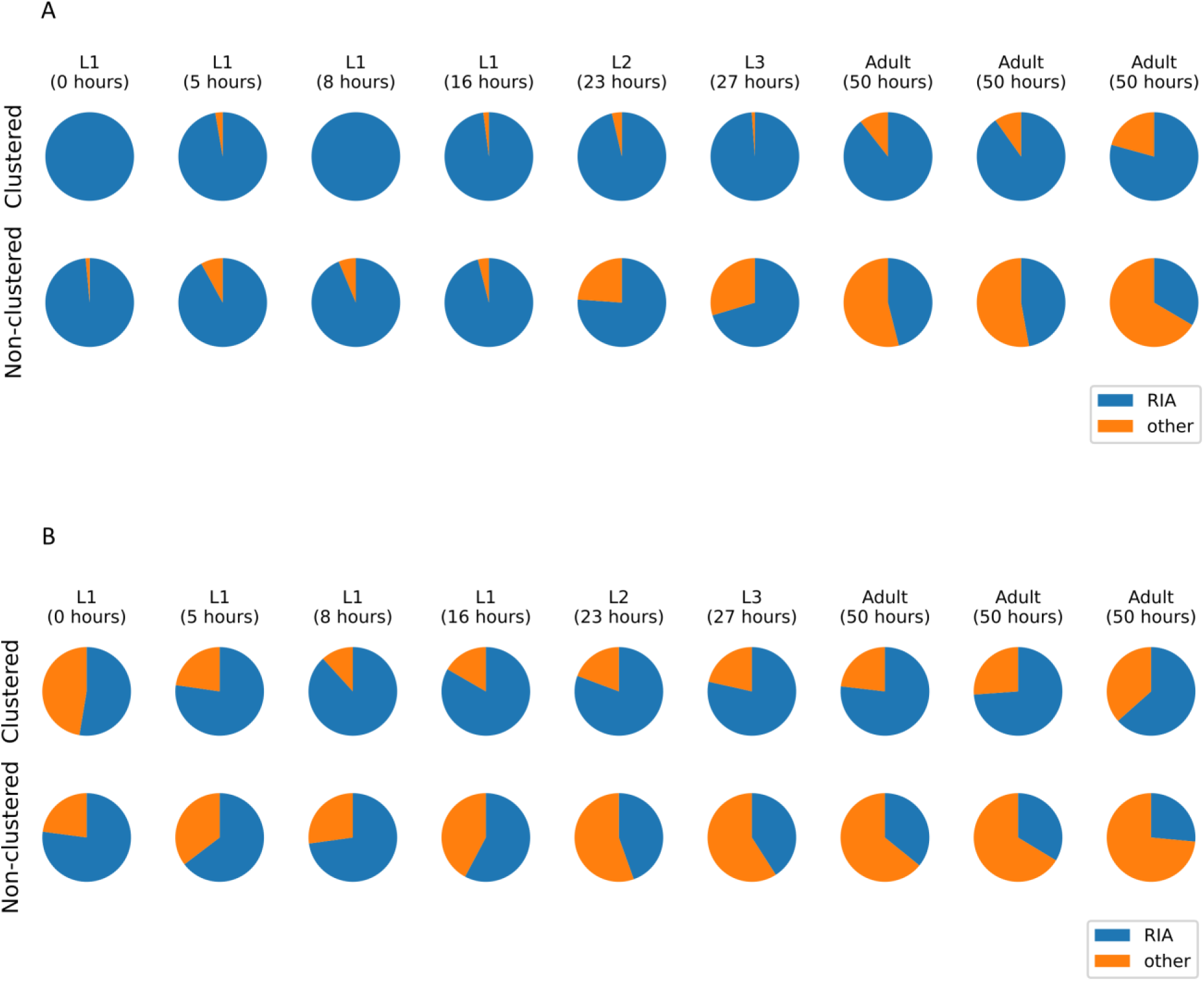
The RIA interneuron emerges as the primary interneuron within the overrepresented tri-neuron circuits to form clustered synapses. (A) Fractions of the overrepresented linear circuits with clustered (top) and non-clustered (bottom) synapses with RIA as the shared connecting interneuron. (B) Fractions of the overrepresented mutually regulating circuits with clustered (top) and non-clustered (bottom) synapses with RIA as the shared pre-synaptic interneuron.

**Supplementary Figure 3.**
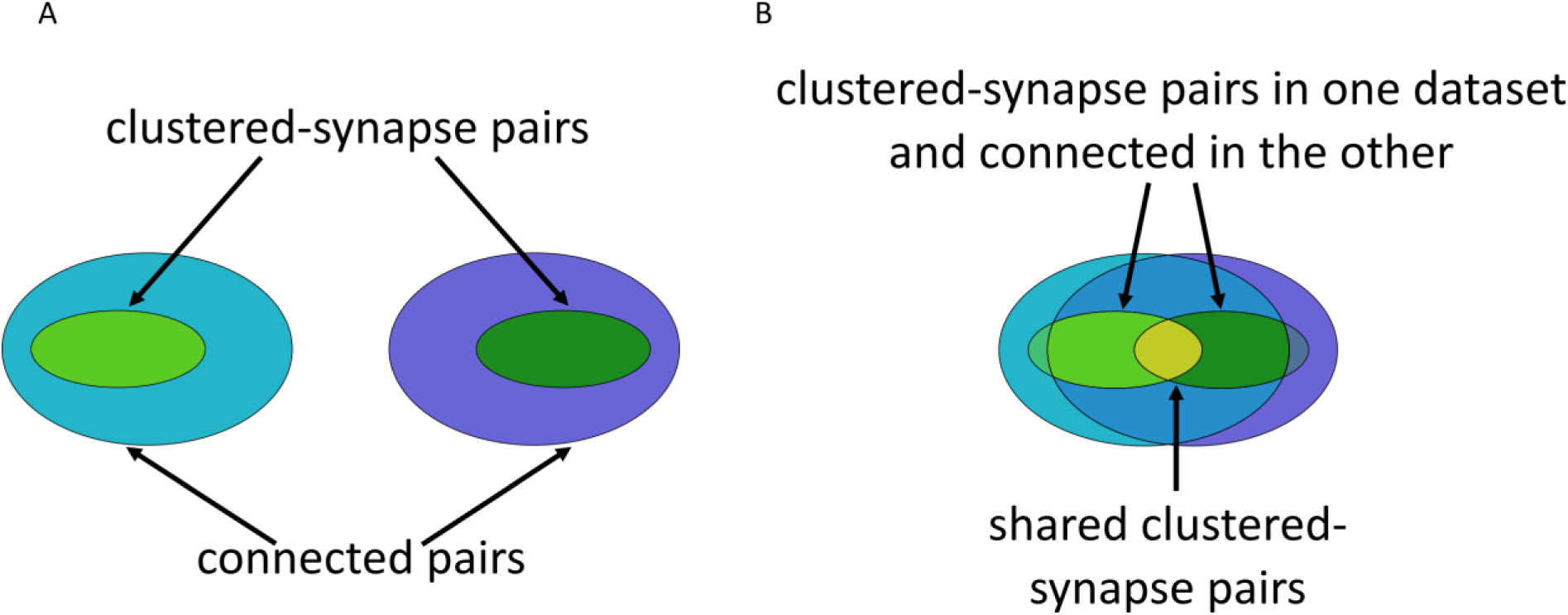
Comparison of neuron pairs with clustered synapses between datasets. (A) Two datasets of connected pair of neurons, each containing a subset of pairs connected by clustered synapses. (B) The number of neuronal pairs with clustered synapses in both datasets (yellow) was divided by the number of pairs with clustered synapses that also appear in the other dataset (the two green sections).

**Supplementary Figure 4.**
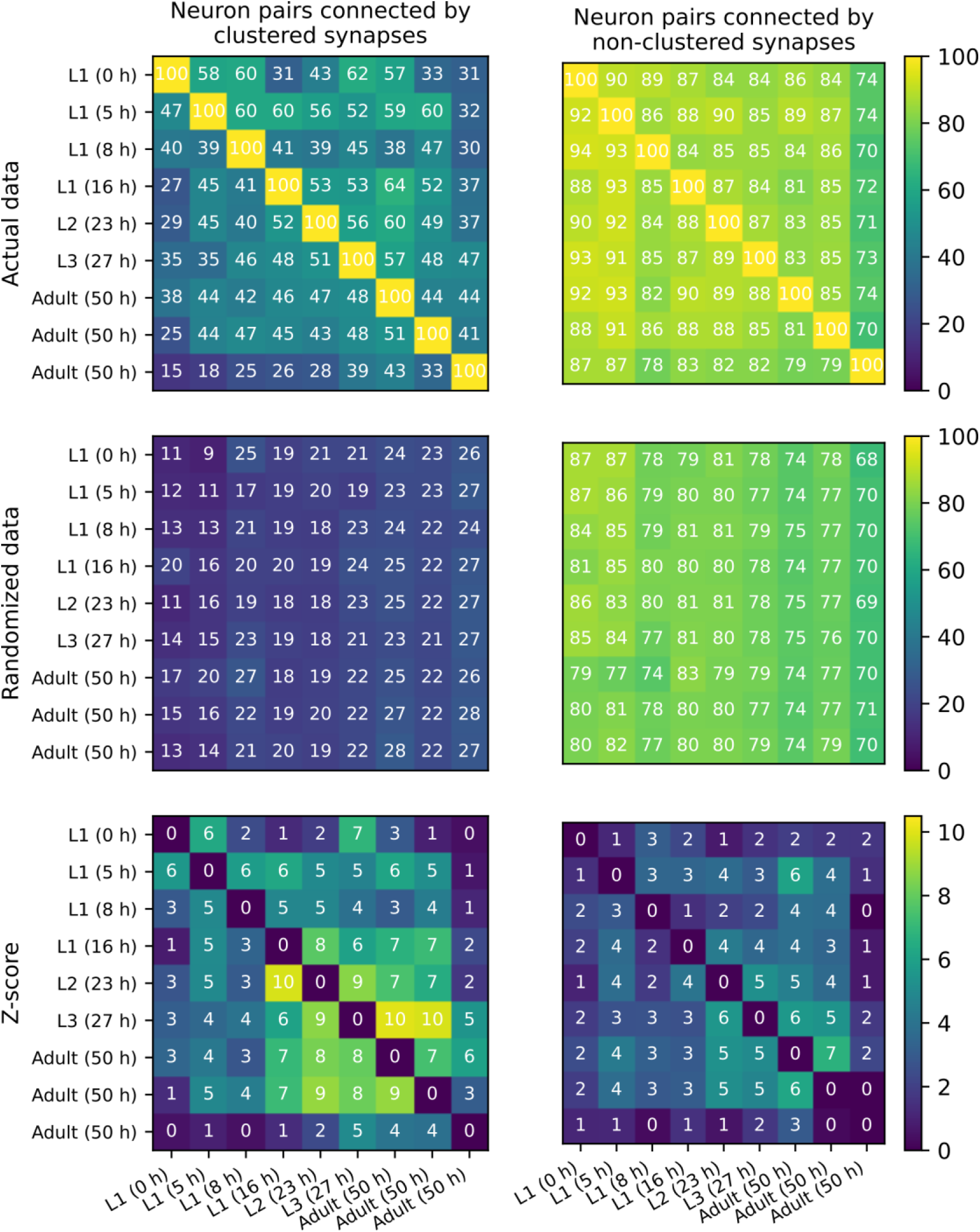
Similarity of neuron pairs connected by at least three synapses. Top: A pairwise analysis indicating the % of neural pairs with clustered synapses in one developmental stage (denoted by the row) that also have clustered synapses in the other developmental stage (denoted by the column). Middle: Repeating the analysis using bootstrap to correct for the number of neural pairs with clustered and non-clustered synapses (see Methods). N=50 iterations. Bottom: Z-scores calculated based on the top and the middle matrices. Left and right panels denote the neural pairs connected by clustered and non-clustered synapses, respectively. The values of the right most and the bottom are for the adult animal connectome compiled by ^17^. All developmental stages as well as the two other adult connectomes are from ^18^.

**Supplementary Figure 5.**
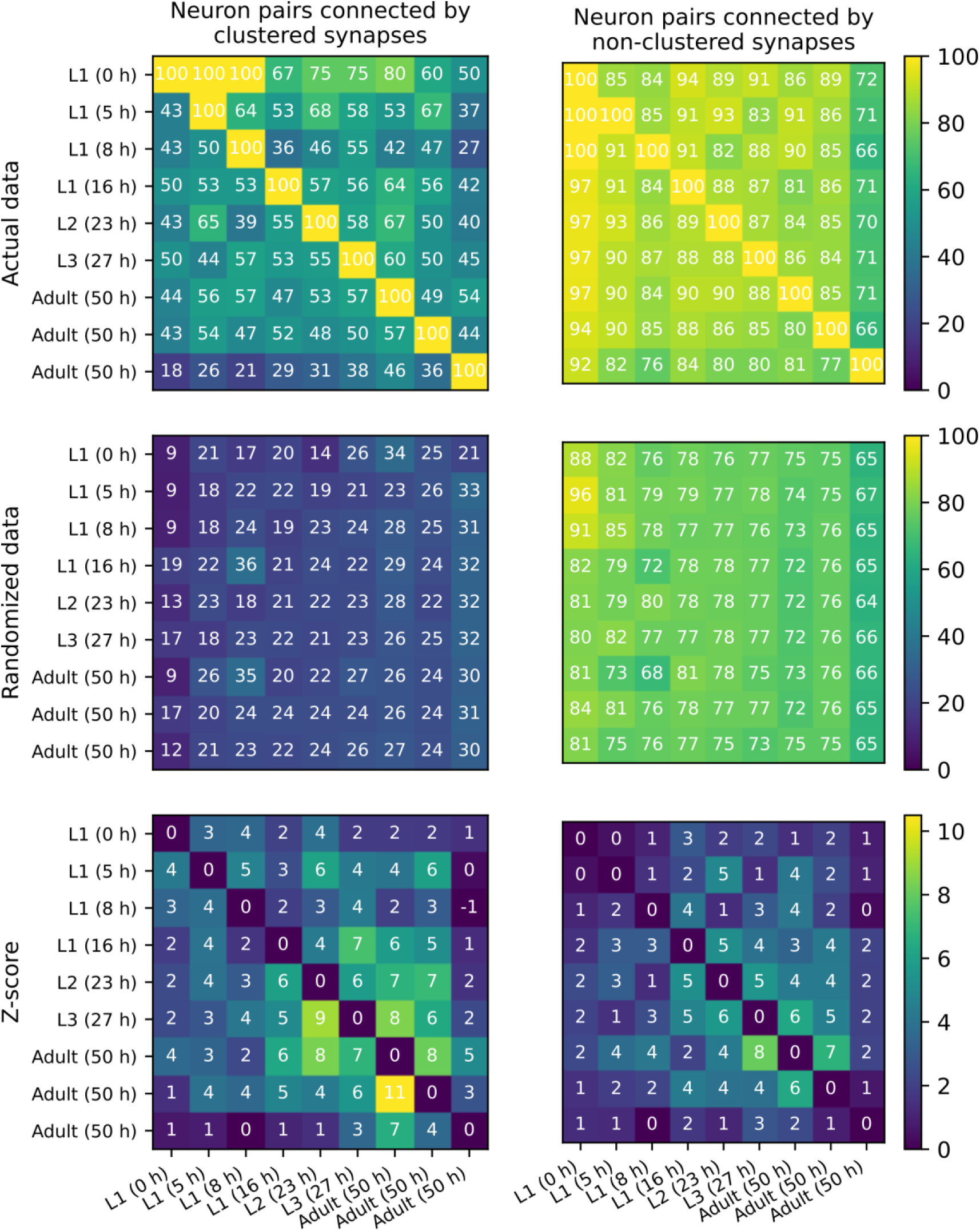
Similarity of neuron pairs connected by at least four synapses. Top: A pairwise analysis indicating the % of neural pairs with clustered synapses in one developmental stage (denoted by the row) that also have clustered synapses in the other developmental stage (denoted by the column). Middle: Repeating the analysis using bootstrap to correct for the number of neural pairs with clustered and non-clustered synapses (see Methods). N=50 iterations. Bottom: Z-scores calculated based on the top and the middle matrices. Left and right panels denote the neural pairs connected by clustered and non-clustered synapses, respectively. The values of the right most and the bottom are for the adult animal connectome compiled by ^17^. All developmental stages as well as the two other adult connectomes are from ^18^.

